# Multiplexed direct detection of barcoded protein reporters on a nanopore array

**DOI:** 10.1101/837542

**Authors:** Nicolas Cardozo, Karen Zhang, Katie Doroschak, Aerilynn Nguyen, Zoheb Siddiqui, Karin Strauss, Luis Ceze, Jeff Nivala

## Abstract

Genetically encoded reporter proteins are a cornerstone of molecular biology. While they are widely used to measure many biological activities, the current number of uniquely addressable reporters that can be used together for one-pot multiplexed tracking is small due to overlapping detection channels such as fluorescence. To address this, we built an expanded library of orthogonally-barcoded Nanopore-addressable protein Tags Engineered as Reporters (NanoporeTERs), which can be read and demuxed by nanopore sensors at the single-molecule level. By adapting a commercially available nanopore sensor array platform typically used for real-time DNA and RNA sequencing (Oxford Nanopore Technologies’ MinION), we show direct detection of NanoporeTER expression levels from unprocessed bacterial culture with no specialized sample preparation. These results lay the foundations for a new class of reporter proteins to enable multiplexed, real-time tracking of gene expression with nascent nanopore sensor technology.

## Main Text

For nearly four decades, reporter proteins have been used as a means to track biological activities such as genetic regulation^1^. Although several different reporter strategies have been developed over this period, the typical number of uniquely addressable reporters that can be used together while sharing a common readout is small^2–5^. This is primarily due to the optical nature of traditional reporters, such as fluorescent protein variants, which have overlapping spectral properties that make simultaneous measurement of unique genetic elements difficult^2^. However, the ability to increase the multiplexability of genetically-encoded protein reporters would enable more comprehensive and scalable monitoring of biological systems, enabling, for instance, high-dimensional phenotyping^6^. This is particularly important for synthetic biology, in which scalable reporter systems are needed to keep pace with the complexity that biological systems can now be engineered in applications such as whole-cell biosensing^7^ and genetic circuit design^8^.

While biomolecular sensing with nanopore sensors has been explored^9^, only recently have high-throughput nanopore sensor platforms emerged for real-time sequencing of DNA^10^ and RNA^11^. The commercial emergence and popularization of these technologies creates an opportunity to build an accessible general nanopore-based platform for direct sensing of engineered reporter proteins. In this context, we present here a new class of genetically-encoded protein reporters, which we term Nanopore-addressable protein Tags Engineered as Reporters (NanoporeTERs, or NTERs), that use commercially-available nanopore sensors (Oxford Nanopore Technologies’ MinION device)^10^ for multiplexed direct protein reporter detection without the need for any other specialized equipment nor laborious sample preparation prior to analysis (**Figures 1a-d**).

**Figure 1:**
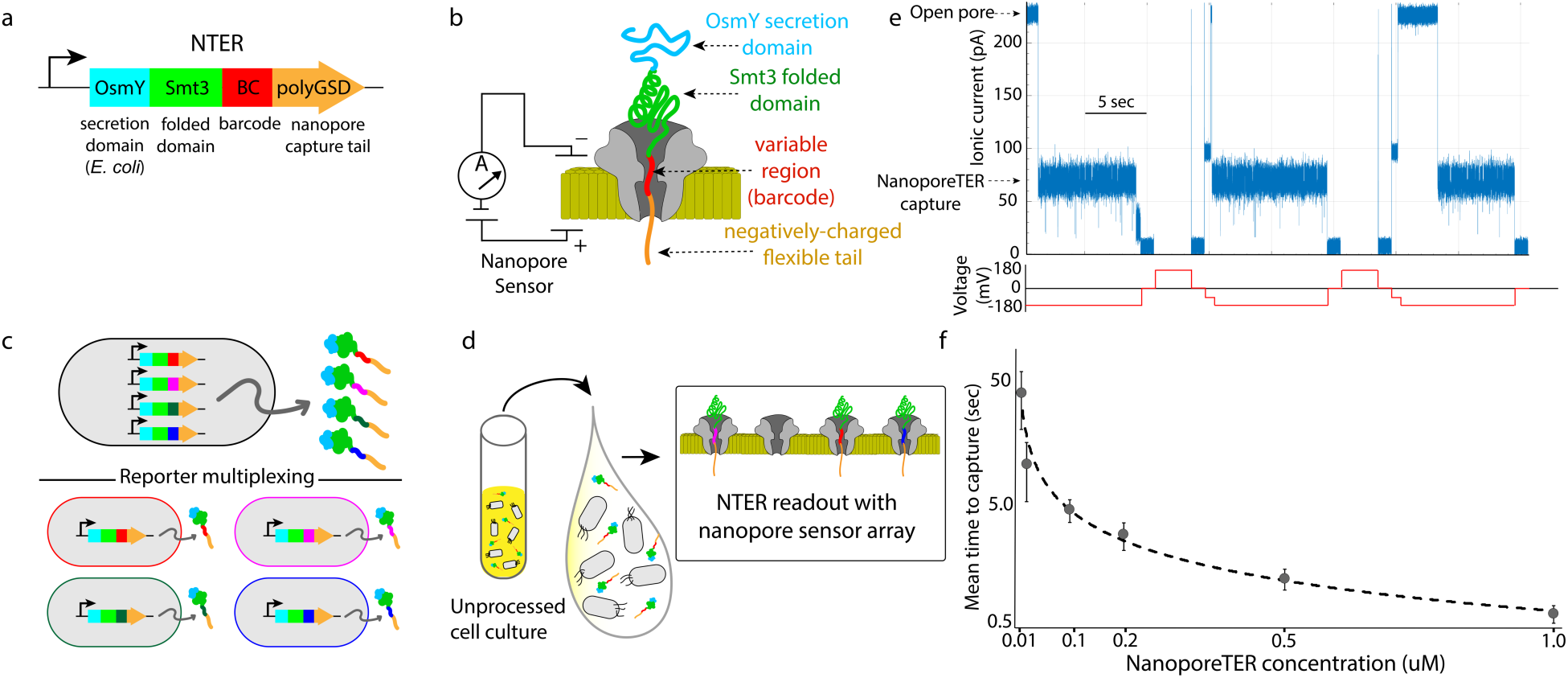
Nanopore-addressable protein Tags Engineered as Reporters. **a**, Gene schematic of NanoporeTER (NTER) design. Cyan: OsmY; promotes extracellular secretion of the reporter protein in *E. coli*. Green: Smt3; folded domain stalls translocation of the protein through the pore and facilitates a “static read” of the NTER barcode within the nanopore sensor. Red: barcode; region of the protein that is held within the sensitive region of the nanopore lumen upon which the changes to the barcode sequence manifest changes to the nanopore ionic current signal. Orange: polyGSD tail; long, flexible, negatively charged C-terminal domain promotes electrophoretic capture of the NTER into the nanopore under an applied voltage. **b**, Schematic of a NanoporeTER captured within a nanopore. **c**, NanoporeTERs are designed to enable multiplexed readout of protein expression, with the potential to report on multiple outputs within a single strain (top), or report on expression across multiple strain types in a one-pot mix (bottom). **d**, Secretion of the NanoporeTERs into the extracellular medium eliminates the need for any sample preparation prior to loading into the nanopore sensor array flow cell. **e**, Example of raw nanopore data generated from a single nanopore showing repeated captures and ejections of NTERY00. **f**, Concentration standard curve showing the relationship between NanoporeTER concentration within a flow cell versus the average time between captures or “reads.”

To develop this method, we started by engineering a protein that is easily detectable by a nanopore sensor and can be expressed and secreted by *E. coli*. We based our initial NTER design on the synthetic protein construct ‘S1’, which we previously developed for unfoldase-mediated nanopore analysis^12,13^. S1 contains a small, folded domain (Smt3) along with a flexible, negatively-charged 65 amino acid C-terminal ‘tail’ composed of glycine, serine, and acidic amino acid residues, in addition to an 11 amino acid ssrA tag^14^. The tail’s lack of structure and net negative charge promotes capture of the protein in a nanopore sensor under an applied voltage. The ssrA tag allows for ClpX-mediated unfolding and translocation of the Smt3 domain, which otherwise inhibits translocation of S1 through the nanopore. For use as a reporter protein in *E. coli*, we modified protein S1 in two ways (**Figure 1a** and **Supp. Figure 1**). First, we replaced the ssrA tag with additional glycine/serine/acidic residues to preserve its nanopore threading activity but prevent targeting of the protein for degradation by ClpXP in-vivo. Second, we added an N-terminal OsmY domain^15^. In *E. coli*, OsmY-tagged proteins are secreted into the extracellular medium^16^. We reasoned secretion would facilitate NTER nanopore analysis by avoiding the need to lyse cells, thereby simultaneously reducing both experimental labor and signal noise that could be generated by non-specific interaction of intracellular molecular species (e.g. DNA, RNA, and other proteins) with the nanopores during analysis. Experiments in BL21 (DE3) *E. coli* showed that expression of this modified version of S1, which we term here ‘NTERY00’, resulted in secretion of the protein into the medium, as detected by SDS-PAGE analysis (**Supp. Figure 2**).

We next purified the secreted NTERY00 by immobilized metal affinity chromatography (IMAC) and then determined if the NTER could be detected on a MinION. To do this, we used an unmodified R9.4.1 flow cell (which uses a variant of the CsgG pore protein^17^) and a custom MinION run script (see Methods). The script applies a constant voltage of −180 mV to all the active pores on the flow cell and statically flips the voltage in the reverse direction in 15 second cycles (ie. 10 seconds ‘ON’ at −180 mV and 5 seconds ‘OFF’ or in ‘Reverse’, see **Figure 1e**). The typical R9.4.1 open pore current level at −180 mV and 500 mM KCl is ∼220 pA. As expected, when NTERY00 was introduced into the flow cell at a concentration of 0.5 uM in these conditions, the current level during each −180 mV portion of the voltage cycle typically underwent a stepwise drop from the open pore value to a consistent lower ionic current state (**Figure 1e** and **Supp. Figure 3**), signaling a putative capture of an NTER within the pore. This current drop was reversible (back to open pore) following reversal of the voltage. We further found that the average time spent in the open pore state before transitioning to the lower ionic current state was NTER concentration dependent (**Figure 1f**). These observations are consistent with a model in which the negatively-charged NTER polyGSD tail is electrophoretically captured in the pore under the applied voltage (−180 mV), and can be ejected from the pore by reversal of the electric field.

If this model is correct, we postulated that the ionic current characteristics of the NTERY00 capture state should be dependent upon the amino acid sequence of the residues residing within the pore’s sensitive limiting constriction. To test this, we made a series of NTER mutants (NTERY01-15) in which a sliding three residue region of the polyGSD sequence was mutated to tyrosines (**Figure 2a**). Tyrosines were chosen because their larger side chain structure was predicted to decrease the ionic current flow through the pore relative to the glycines and serines of NTERY00 when captured within the pore. Following purification and MinION analysis of NTERs 01-15, we found the capture state to be NTER mutant-dependent up to NTERY08, after which we observed NTER mutants 09-15 to have signal characteristics indistinguishable from NTERY00 (**Figures 2b, c, and d**, and **Supp. Figures 3 and 4**). These results support a model in which the first ∼17 amino acids of the polyGSD tail reside with the CsgG nanopore’s sensitive region and contribute to its ionic current signature during a capture event. This also sets an upper bound to the number of possible NTER barcode sequences at 20^17^ (∼10^22^).

**Figure 2:**
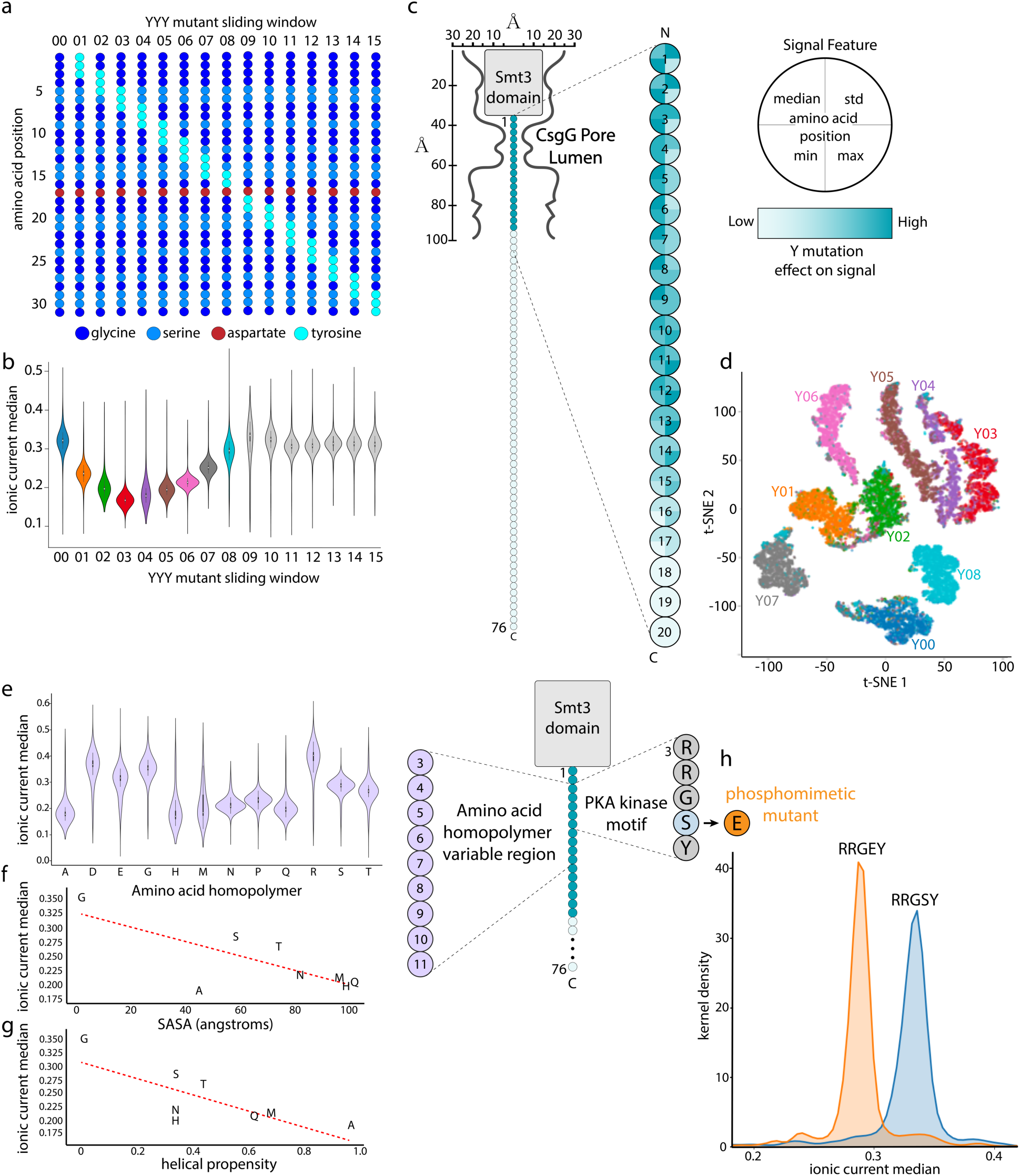
Mapping the NanoporeTER sequence and nanopore signal space on a MinION. **a**, Schematic of the NTERs Y00-15 mutant sequences in which a sliding block of three tyrosine mutations was introduced along the NanoporeTER polyGSD barcode and tail region to map the NTER’s nanopore-sensitive region and define the potential barcode sequence space. **b**, Violin plot showing the median ionic current level (normalized to the open pore level) of the nanopore capture state for NTERs Y00-15. The introduction of the three tyrosine block (YYY), reduces the ionic current level in a position-dependent manner for positions 01-08. The median current level returns to the baseline (NTER Y00) level starting at position 9 and through position 15, supporting a model in which the first 17 amino acids of the polyGSD tail contribute to the observed NTER ionic current signature, and defining the NTER barcode region. Each NTER distribution is composed of several thousand single-molecule measurements. **c**, Simple structural model of the NTER position within the nanopore during a read (capture event). A heat map displaying the relative change to specific signal features (median, standard deviation, minimum, and maximum) is projected onto the NTER tail residue positions (1-20) that were mutated in NTERs Y00-15, showing the relative magnitude of effect tyrosine mutations at each residue have on the NTER’s nanopore ionic current signal. **d**, t-SNE plot clustering NTER reads (each read is represented as a single point) based on ionic current signal features (mean, std, min, max, median), and colored by the NTER’s barcode identity (Y00-08). n = ∼4000 events per barcode class. **e**, Violin plot showing the median ionic current level (normalized to the open pore level) of the nanopore capture state for amino acid homopolymer NTERs alanine (A), aspartate (D), glutamate (E), glycine (G), histidine (H), methionine (M), asparagine (N), proline (P), glutamine (Q), arginine (R), serine (S), and threonine (T). Each NTER distribution is composed of ∼1500 single-molecule measurements. **f**, Scatter plot showing the relationship between amino acid solvent accessible surface area (SASA) versus the respective amino acid homopolymer NTER mutant’s median ionic current level (normalized to the open pore level). **g**, Scatter plot showing the relationship between amino acid helical propensity versus the respective amino acid homopolymer NTER mutant’s median ionic current level (normalized to the open pore level). **h**, Kernel density plot comparing the ionic current median (normalized to the open pore level) of reads generated by an NTER containing a PKA phosphorylation motif (RRGSY) within its barcode region to those with a phosphomimetic mutation (RRGEY). Each NTER distribution is composed of several thousand single-molecule measurements.

After determining the number of amino acids that contribute to the NTER nanopore signal (the NTER sequence space), we next sought to determine how different amino acid types modulate the ionic current through the pore. These results help define the possible future NTER signal space. To investigate this, we constructed NTER variants in which positions 3-11 within the polyGSD region were mutated to all the 20 possible standard amino acid homopolymers. **Figure 2e** and **Supp. Figure 5** show the signal features of the ionic current levels for 12 out of the 20 NTER homopolymer mutants (the homopolymers C, F, I, K, L, V, W, and Y, most of which have significant hydrophobic character, did not express sufficient soluble protein). To see how the different amino acid physical properties contribute to the NTER ionic current, we investigated whether certain properties correlate with different signal features. While no strong correlations were found across all the 12 amino acid types, we did find that the median current level moderately correlated with both amino acid volume and helical propensity within the uncharged amino acid types (R = ∼0.75 for each, **Figure 2f,g**).

Next, to probe the potential of this method to resolve between amino acid barcodes with subtler sequence differences (for example, point mutations or post-translational modifications), we cloned and tested two additional NTER barcodes based on the protein kinase A (PKA) phosphorylation motif^18^. The first PKA-based barcode contained a canonical PKA motif (RRGSY), while the second had a single amino acid difference (RRGEY) that mimics the PKA motif’s phosphorylated serine state in structure and charge (commonly referred to as a ‘phosphomimetic’, **Figure 2h**). Following purification and MinION analysis of these two NTERs, we found that the phosphomimetic barcode could be distinguished from the canonical PKA motif barcode, as the two barcodes typically had substantially different nanopore ionic current state medians (**Figure 2h**). These results suggest the potential of using NanoporeTERs to report on the activity of enzymes that regulate specific post-translational modifications, such as phosphorylation and methylation.

Finally, having explored the potential NTER barcode sequence space, signal space, and sensitivity to single residue modifications, we sought to demonstrate proof-of-principle NTER applications for multiplexed tracking of gene expression. To do this, we first used supervised machine learning to train classifiers that could accurately discriminate amongst combinations of the NTER barcodes explored above. Using either a set of engineered signal features as input to a Random Forest (RF) classifier or the raw ionic current signal directly into a Convolutional Neural Network (CNN) (**Figure 3a**), we used our purified NTER datasets for model training and validation. Both models achieved similar accuracies that ranged from ∼80-90% depending on the model hyperparameters and barcode set (**Figure 3b**, see Methods).

**Figure 3:**
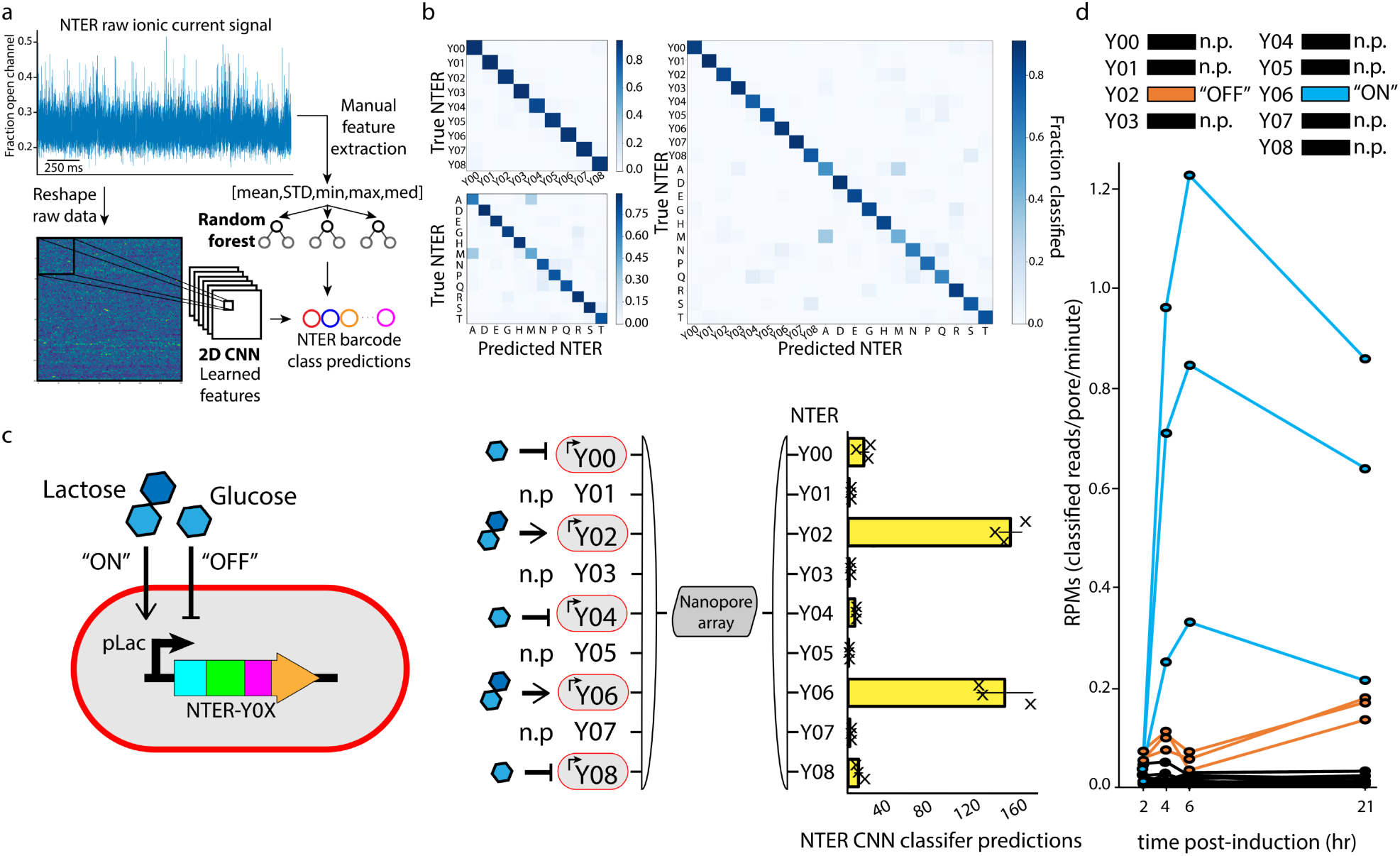
Classification and multiplexed detection of NanoporeTER expression levels with a MinION. **a**, Raw ionic current data was classified using either a set of engineered features (mean, std, min, max, and median) or the unprocessed signal directly, and input into either a Random Forest or Convolutional Neural Network classifier, respectively. **b**, Confusion matrices showing the Random Forest test set classification accuracies on models using different combination of NTER barcodes. Top left: NTERs Y00-08. Bottom left: amino acid homopolymer mutants A, D, E, G, H, M, N, P, Q, R, S, and T. Right: Both the NTERs Y00-08 and amino acid homopolymer mutants. **c**, Schematic showing the gene construct used for controllable NTER expression. Lactose is used to induce NTER expression (“ON”), while glucose inhibits expression (“OFF”). The diagram and bar plot on the right shows the results of a mixed culture experiments in which NTER expression was induced for NTERs Y02 and Y04, and inhibited for NTERs Y00, Y02, and Y08. NTERs Y01, Y03, Y05, and Y07 were held out of the experiment as negative controls. Plot shows the total number of reads classified as each NTER barcode during MinION analysis. **d**, Line plot showing a time course of NTER expression levels as determined by the rate of classified reads (reads/pore/min) for each NTER barcode. NTER Y06 was induced, while NTER Y02 was inhibited. The other NTERs were held out as negative controls and show false-positive classification rates. Three replicates for each condition are plotted.

We then used the best performing CNN that was trained on NTERs Y00-08 (in addition to a background/noise class, see Methods) to determine the relative NTER expression levels within bacterial cultures composed of mixed populations of strains engineered with different NTER-tagged plasmid-based circuits. To do this, we grew independent mono-barcoded cultures overnight with NTER expression either induced or inhibited (by autoinduction media containing lactose or LB supplemented with glucose, respectively). In the morning, just prior to nanopore readout, the cultures were mixed into a single solution and diluted into MinION running buffer and loaded directly into a flow cell for analysis. Importantly, these cell cultures underwent no processing or purification prior to analysis, in contrast to our previous experiments. Results from these experiments showed higher classification counts for the NTER barcodes for which expression was induced (NTERs 02 and 06), and lower counts for strains that were inhibited (glucose: NTERs Y00, Y04 and Y08) or not present at all in the mixed population (NTERs Y01, Y03, Y05, and Y07) for all replicates (**Figure 3c**) over a ten minute MinION runtime. We then conducted a time course experiment in which we tracked expression of two different NTERs over multiple hours, one of which was induced with IPTG (NTERY06), and the other of which NTER expression was inhibited with glucose (NTERY02). Again, cultures were grown independently, but then mixed just prior to nanopore readout. **Figure 3d** shows the results of this time course (and replicates) during ten minutes of MinION analysis at 2, 4, 6, and 21-hour timepoints following induction (NTERY06) or inhibition (NTERY02) of the NanoporeTER circuit. Again, the rate of NTER classification (reads/pore-minute) was substantially higher for the induced NTERY06 circuits, compared to the uninduced NTERY02 circuits. Importantly, leaky expression of NTERY02 was still detectable over the background false-positive classification rates for the NTER barcodes that were not present at all in the experiment (Y00, Y01, Y03, Y04, Y05, Y07 and Y08). These results demonstrate that NanoporeTERs can be used as reliable reporters of relative protein expression levels.

In conclusion, we have laid the foundations for a new class of multiplexable protein reporters (NanoporeTERs) that can be analyzed using a commercially available nanopore sensor array, the ONT MinION. While we have characterized here a set ∼20 orthogonal NanoporeTERs, we are confident that this number can be increased significantly with the following strategies: 1) high-throughput methods to empirically characterize more barcode sequences for classifier training, 2) engineering NanoporeTERs to contain multiple barcode regions that can be consecutively read out with the aid of processive motor proteins^12,13^ or voltage-mediated translocation^19^, which would allow the number of orthogonal NTERs to scale exponentially with the number of individually characterized barcodes, and 3) semi-supervised machine learning models trained to accurately predict the sequence of empirically uncharacterized NTER barcodes given only their nanopore signal^20^. NanoporeTERs should also not be limited to use in *E. coli*, as their modular design requires that only the secretion domain be modified, of which many different N-terminal secretion domains have been characterized in a range of diverse organisms^21–23^.

We foresee many potential NanoporeTER applications, including simultaneously reading the protein-level outputs of many genetically engineered circuit components in one-pot, enabling more efficient debugging and tuning than current analysis methods. For instance, in comparison to traditional sets of fluorescent protein reporters, NanoporeTERs have a (potentially much) larger sequence and signal space that allows for the simultaneous analysis of a greater number of unique genetic elements in a single experiment (multiplexing). And while RNA-seq is another strategy that can be used to measure the transcriptional output of many circuits in parallel with high-throughput DNA sequencing technology^24^, our method has the advantages of 1) little to no sample preparation, which makes it more amenable to automation^25–27^ and reduces both time to analysis (latency) and cost, and 2) direct detection of outputs at the protein level. The latter advantage opens new opportunities to custom engineer reporters with NTER barcodes that can report on both protein expression and specific post-translational modifications simultaneously. We anticipate this capability will be especially useful as the nascent field of synthetic protein-level circuit engineering advances^28^.

## Acknowledgments

We thank additional members of the Molecular Information Systems Lab for helpful discussion and feedback on this work. The OsmY expression plasmid was generously provided by Cassie Bryan and Lauren Carter (UW). We also thank Andy Heron and Rich Gutierrez (ONT) for providing the configurable MinION run script and discussions on its use, and Miten Jain (UCSC) for a custom Matlab script that facilitated visualization of the raw MinION data.

## Author contributions

N.C., K.Z., and A.N. performed wet lab experiments. K.Z. and K.D. developed the data analysis pipeline and performed computational analyses. Z.S. implemented the machine learning approach. K.S., L.C., and J.N. supervised the project. J.N. conceived and directed the project. All authors contributed to writing and editing of the manuscript.

## Data and materials availability

Data and code are available upon request and on github.com/uwmisl/NanoporeTERs.

## Supplementary Materials

Materials and Methods

Figures S1-S5

