## Supplementary Materials for "Multiplexed direct detection of barcoded protein reporters on a nanopore array"

**This PDF file includes:**

Materials and Methods
Figs. S1 to S5

### Materials and Methods

NanoporeTER construction, cloning, expression, and purification

The initial NanoporeTER protein was constructed with a gBlock (Integrated DNA Technologies) composed of the Smt3 and tail sequence and cloned into plasmid pCDB180 downstream of the OsmY domain. The Q5 site-directed mutagenesis method (New England Biolabs) was used to generate the different NTER barcode mutants. All cloning was performed using the 5-alpha competent *E. coli* strain following NEB's cloning protocol (New England Biolabs). Sequence verification was obtained through Genewiz Inc. Expression of the NanoporeTER protein was done in BL21 (DE3) *E. coli* strain using Overnight Express instant TB medium (Novagen).

Proteins were purified via immobilized metal affinity chromatography (IMAC) using TALON metal affinity cobalt resin (Takara). The purification used the associated buffer set from Takara, following their specified protocol. Proteins were concentrated using Amicon Ultra 0.5 mL centrifugal filters with Ultracel 30K (Amicon). The final concentration of proteins averaged ~7 mg/ml from 5 mL overnight cultures. The purified proteins were stored for long-term storage at -80C in 10 uL aliquots, as well as for short-term storage at 4C.

Raw culture mixing experiments

Cultures were picked from single colonies on plates and used to inoculate 3mL LB supplemented with 0.5 mM IPTG and kanamycin (induced), or 3mL LB supplemented with 0.2% glucose and kanamycin (inhibited). After overnight incubation at 37C with shaking, cultures were equally mixed together in a total volume of 45uL, 50uL 4X C17 buffer (2 M KCl, 100 mM HEPES, pH 8), and 105 uL water (total volume 200uL). This solution was then immediately loaded into a MinION flow cell for analysis.

Time course

Time course experiments were performed by diluting 30uL of overnight cultures (LB) into 3mL fresh LB supplemented with 0.5 mM IPTG and kanamycin (induced), or 3mL fresh LB supplemented with 0.2% glucose and kanamycin (inhibited). The cultures were placed in a shaker/incubator at 37C to allow for culture growth. Samples were then collected at 2, 4, 6, and 21-hour time-points. At each time-point, cultures were equally mixed together in a total volume of 10 uL, 50uL 4X C17 buffer, and 140 uL water (total volume 200uL). This solution was then immediately loaded into a MinION flow cell for analysis.

MinION experiments

All experiments were performed with unmodified R9.4.1 MinION flow cells (Oxford Nanopore Technologies) by diluting analyte solution into C17 buffer for a final concentration of 0.5M KCl and 25mM HEPES (pH 8), into the flow cell priming port. Flow cells were run on the MinION at a temperature of 30°C and a run voltage of -180mV with a 10khz sampling frequency and 15 second static flip frequency. Use of a modifiable MinKNOW script (available from ONT) enabled voltage flipping cycle parameters to be set as well as collection of raw current data across the entire run. Individual flow cells could be reused for different analytes after flushing them with 1mL C17 buffer three times between experiments. Flow cells were stored at 4°C in C18 buffer (150mM potassium ferrocyanide, 150mM potassium ferricyanide, 25mM potassium phosphate, pH8) when not in use.

Nanopore Signal Analysis, Quantification, and Classification

The analysis pipeline for a NanoporeTER sequencing run begins with extracting the segments of the raw nanopore signal that contain capture events. A capture is defined as a region where the signal current falls below 70% of the open pore current for a duration of at least one millisecond. The fractional current values (as compared to open pore current) computed from the segmentation process, as well as the start and end times of each capture, are saved in separate data files. This information is then passed through a general filter that separates putative NanoporeTER captures from noise captures based on features of the normalized raw current (mean, standard deviation, minimum, maximum, median) as well as the duration of the capture. Captures that pass this initial filter are then fed into a classifier and classified as a specific NTER barcode or a background/noise blockade. The metadata for the captures within each NTER class are subsequently fed to a quantifier which calculates the average time elapsed between those captures and converts this time to the predicted NTER concentration using a standard curve. An alternative method of quantification is to calculate the number of reads per class per active pore per minute (reads/pore-minute or RPMs). In addition to the NTER data sets, a background/noise class data set was also used in training the models to recognize data generated from non-NTER-specific pore blockages that made it through the filtering step. This data was collected from experiments in which only running buffer, LB media, or NTER-free *E. coli* cultures were loaded into the flow cell.

We explored two different classifiers for NTER barcode discrimination. The first, a Random Forest model, was implemented in scikit-learn (sklearn.ensemble.RandomForestClassifier). The second classifier was a CNN implemented in PyTorch. An 80/20 train/test split was used to generate the classification accuracy estimates and confusion matrix results. For both models, only the first two seconds of each capture were considered for analysis. The Random Forest was trained on an array composed of the mean, standard deviation, minimum, maximum, and median of that two second window. Default Random Forest hyperparameters were modified to: n_estimators=300 and max_depth=100. The CNN used the two seconds of raw signal directly as input following reshaping of the 1D signal into a 2D structure. The neural network was composed of four 2D convolutional layers each with ReLU activation and max pooling. These were followed by a fully connected layer which had a log-sigmoid activation function, and then a final output layer of the same size as the number of NTER classes considered in the experiment. Full model details and code can be found at https://github.com/uwmisl/NanoporeTERs.

NTER amino acid sequence:

MTMTRLKISKTLLAVMLTSAVATGSAYAENNAQTTNESAGQKVDSSMNKVGNFMDDSAITAKVKAALVDHDNIKSTDISVKTDQKVVT
LSGFVESQAQAEEAVKVAKGVEGVTSVSDKLHVRDAKEGSVKGYAGDTATTSEIKAKLLADDIVPSRHVKVETTDGVVQLSGTVDSQ
AQSDRAESIAKAVDGVKSVKNDLKTKMGHHHHHHHHHHGSLQDSEVNQEAKPEVKPEVKPETHINLKVSDGSSEIFFKIKKTTPLRRL
MEAFAKRQGKEMDSLRFLYDGIRIQADQAPEDLDMEDNDIIEAHREQIXXXXXXXXXXXXXXXXXXXXXXXXXXXXXXDGGSSGGS
GGDGSSGDGGSDGDSDGSDGDGDSDGDDGGDDEDDGSDD*

```
NTERY00 X = GGGGSSGGSGGSGSSGDGGSSGGSGGSGSSG
```

YYY mapping mutants:

```
NTERY01 X = YYYGSSGGSGGSGSSGDGGSSGGSGGSGSSG
NTERY02 X = GGYYYSGGSGGSGSSGDGGSSGGSGGSGSSG
NTERY03 X = GGGGYYYGSGGSGSSGDGGSSGGSGGSGSSG
NTERY04 X = GGGGSSYYYGGSGSSGDGGSSGGSGGSGSSG
NTERY05 X = GGGGSSGGYYYSGSSGDGGSSGGSGGSGSSG
NTERY06 X = GGGGSSGGSGYYYSSGDGGSSGGSGGSGSSG
NTERY07 X = GGGGSSGGSGGSYYYGDGGSSGGSGGSGSSG
NTERY08 X = GGGGSSGGSGGSGSYYDGGSSGGSGGSGSSG
NTERY09 X = GGGGSSGGSGGSGSSGDYYSSGGSGGSGSSG
NTERY10 X = GGGGSSGGSGGSGSSGDGYYYGGSGGSGSSG
NTERY11 X = GGGGSSGGSGGSGSSGDGGSYYYSGGSGSSG
NTERY12 X = GGGGSSGGSGGSGSSGDGGSSGYYYGSGSSG
NTERY13 X = GGGGSSGGSGGSGSSGDGGSSGGSYYYGSSG
NTERY14 X = GGGGSSGGSGGSGSSGDGGSSGGSGGYYYSG
NTERY15 X = GGGGSSGGSGGSGSSGDGGSSGGSGGSGYYY
```

Homopolymer mutants:

```
NTER A X = GGAAAAAAAAASGSSGDGGSSGGSGGSGSSG
NTER D X = GGDDDDDDDDDSGSSGDGGSSGGSGGSGSSG
NTER E X = GGEEEEEEEEESGSSGDGGSSGGSGGSGSSG
NTER G X = GGGGGGGGGGGSGSSGDGGSSGGSGGSGSSG
NTER H X = GGHHHHHHHHHSGSSGDGGSSGGSGGSGSSG
NTER M X = GGMMMMMMMMMSGSSGDGGSSGGSGGSGSSG
NTER N X = GGNNNNNNNNNSGSSGDGGSSGGSGGSGSSG
NTER P X = GGPPPPPPPPPSGSSGDGGSSGGSGGSGSSG
NTER Q X = GGQQQQQQQQQSGSSGDGGSSGGSGGSGSSG
NTER R X = GGRRRRRRRRRSGSSGDGGSSGGSGGSGSSG
NTER S X = GGSSSSSSSSSSGSSGDGGSSGGSGGSGSSG
NTER T X = GGTTTTTTTTTSGSSGDGGSSGGSGGSGSSG
```

PKA motif mutants:

```
NTER PKA                 X = GGRRGSYYSGGSGSSGDGGSSGGSGGSGSSG
NTER PKA_phosphomimetic X = GGRRGEYYSGGSGSSGDGGSSGGSGGSGSSG
```

**Supplementary Figure 1** NanoporeTER sequences used in this work. Blue: OsmY domain. Black: His-tag. Green: Smt3 domain. Red: Variable region (contains barcode). Orange: PolyGSD tail.

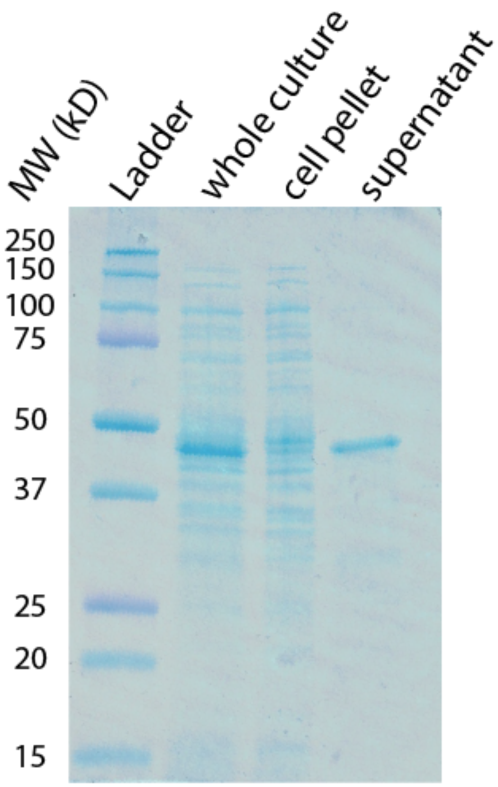


**Supplementary Figure 2** NanoporeTERs are secreted into the extracellular medium. SDS-PAGE analysis of overnight culture of an E. coli strain transformed with a plasmid expressing NTER00 (expected MW is 40.2 kilodaltons). Lanes: 1, Ladder. 2, raw whole culture (cells and growth medium). 3, cell pellet resuspended in water following centrifugation. 4, Growth medium supernatant following centrifugation.

**Supplementary Figure 3** Representative MinION nanopore ionic traces of NTER barcodes analyzed in this work. Three consecutive "reads" are included in each trace.

NTER00

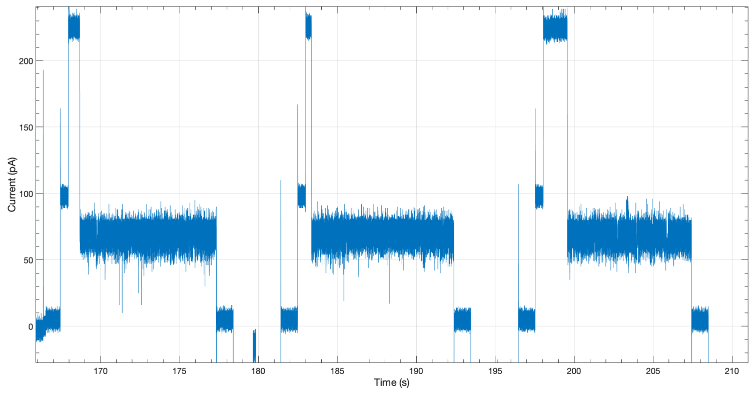


NTER01

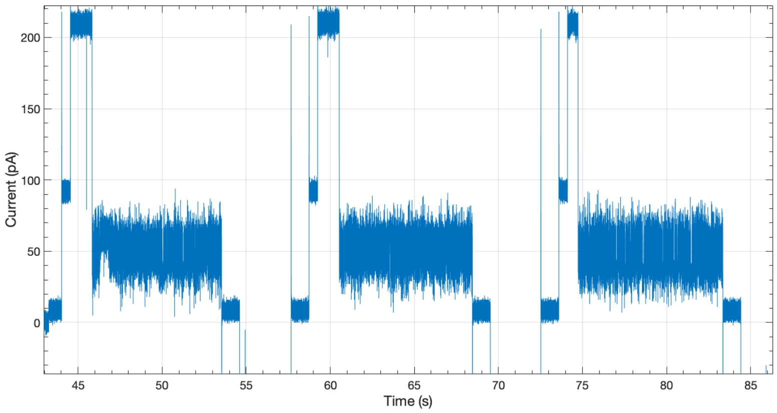


NTER02

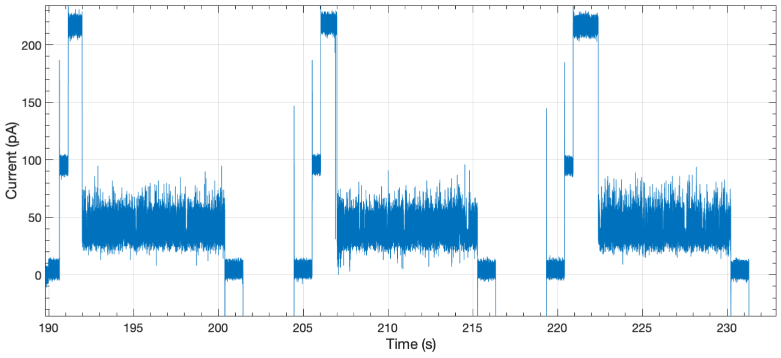

**Supplementary Figure 3 (cont.)**

NTER03

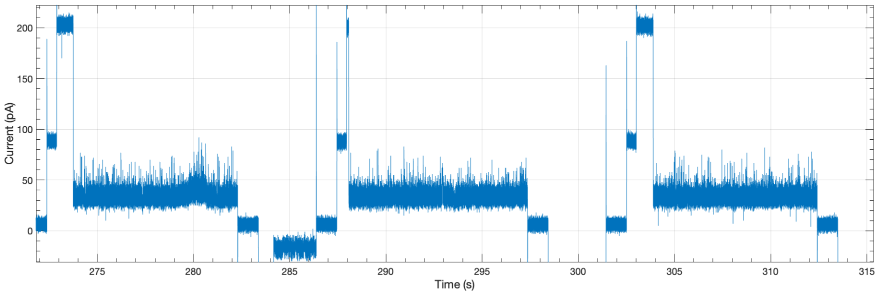


NTER04

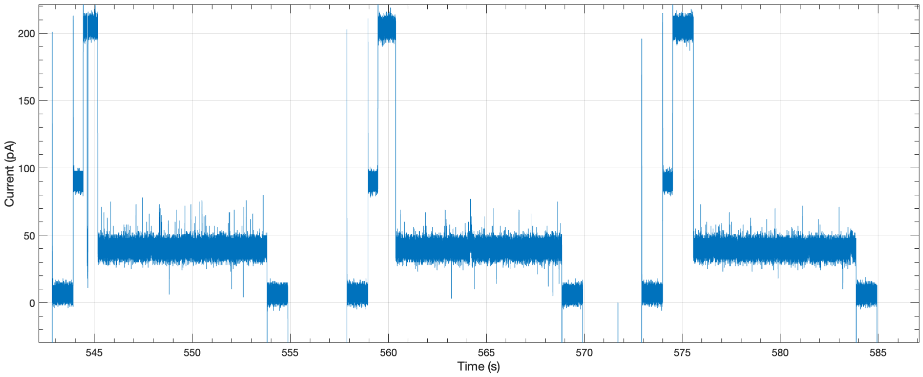


NTER05

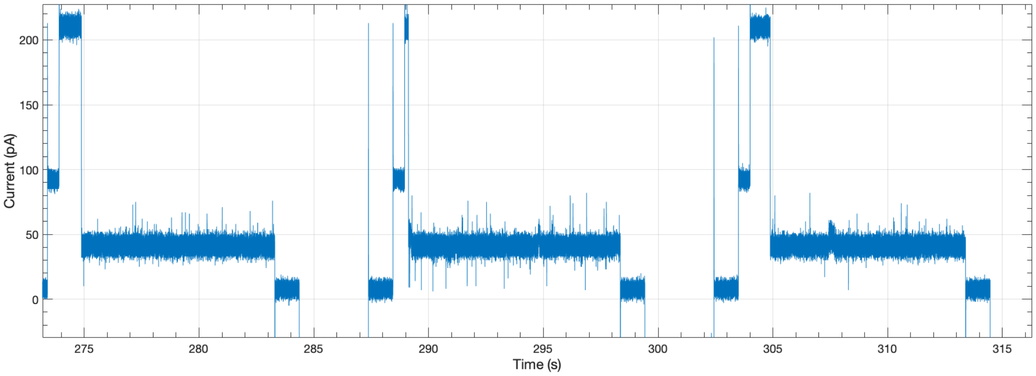

**Supplementary Figure 3  (cont.)**

NTER06

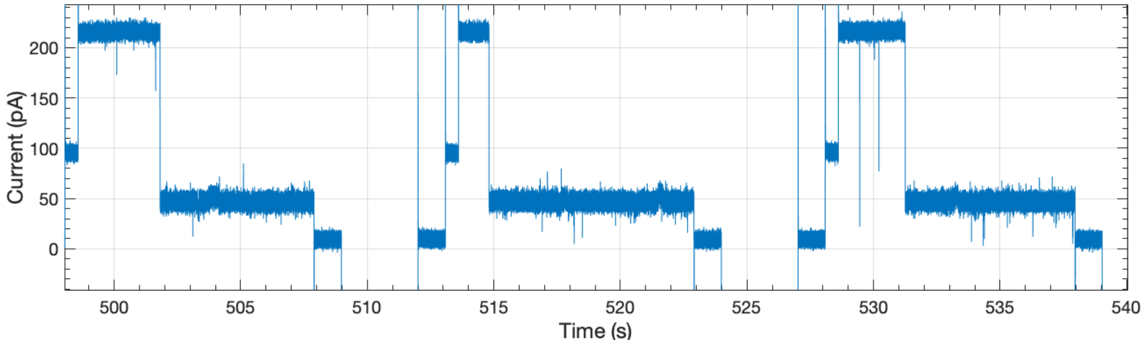


NTER07

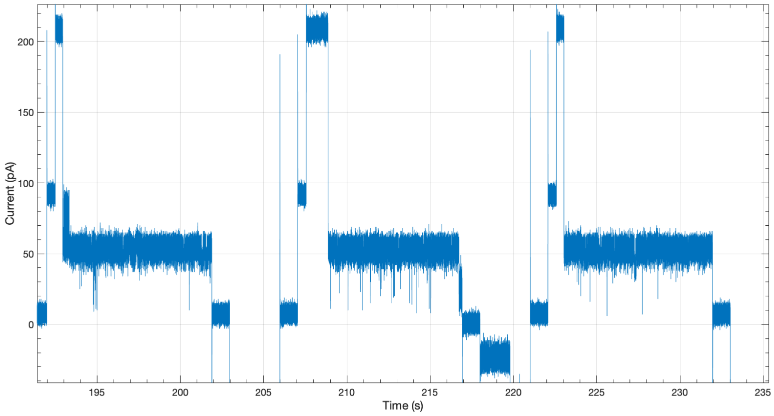


NTER08

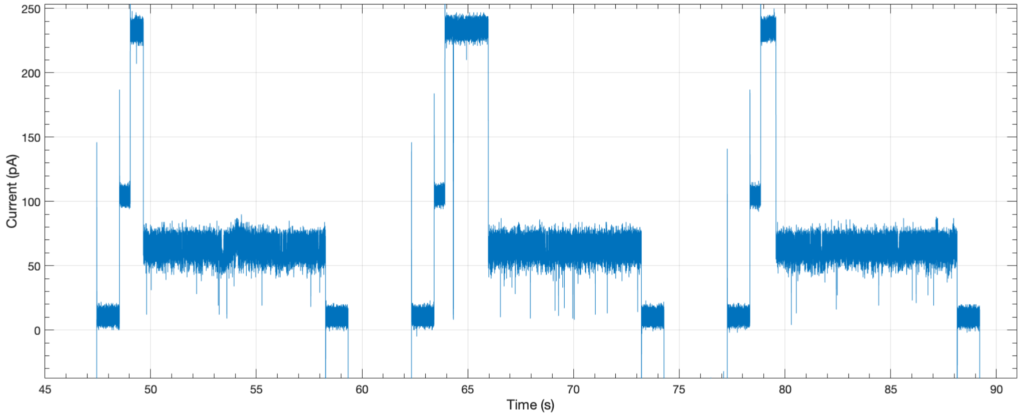

**Supplementary Figure 3 (cont.)**

NTER homopolymer A (Alanine)

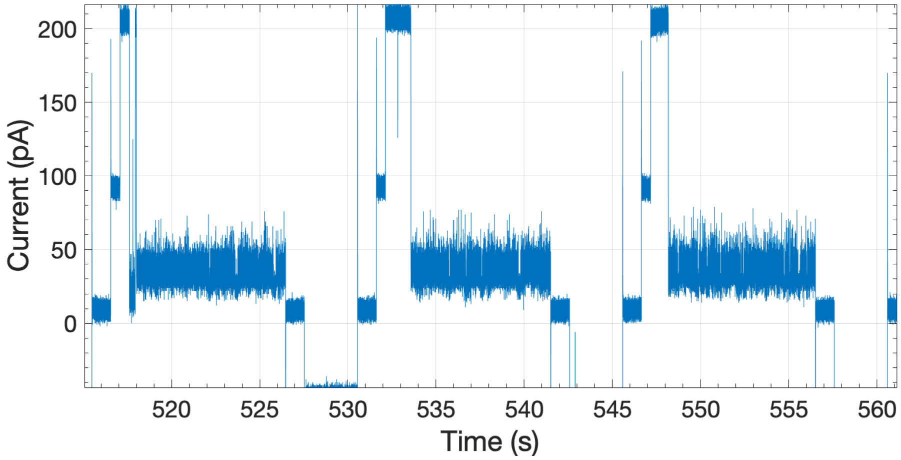


NTER homopolymer R (Arginine)

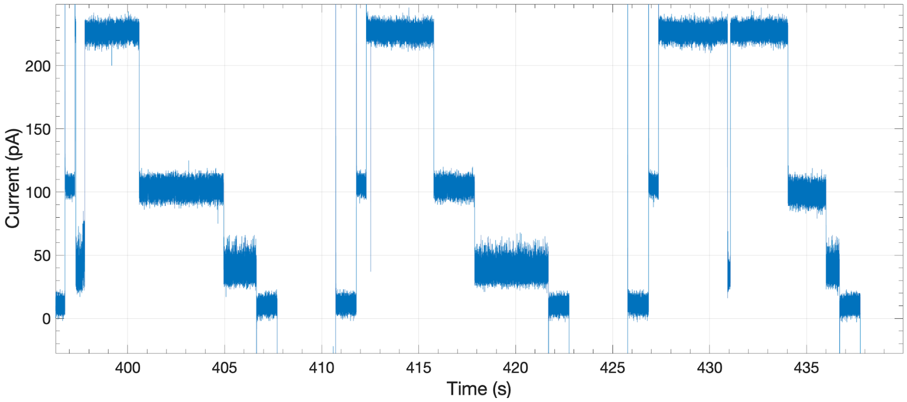


NTER homopolymer N (Asparagine)

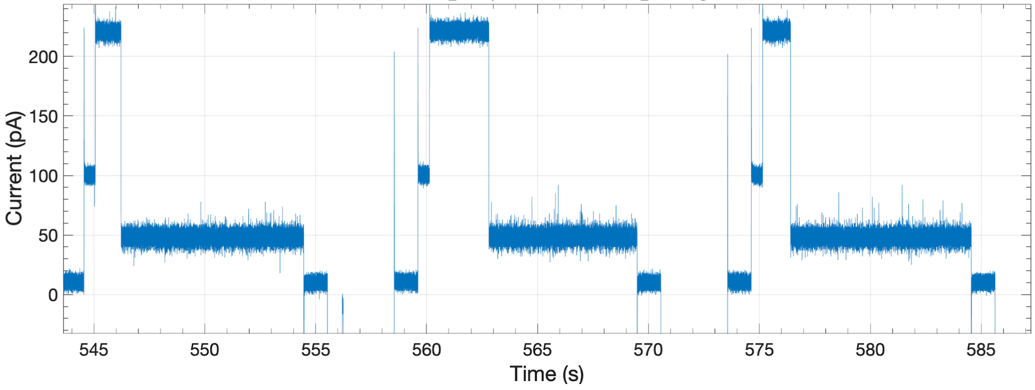

**Supplementary Figure 3 (cont.)**

NTER homopolymer D (Aspartate)

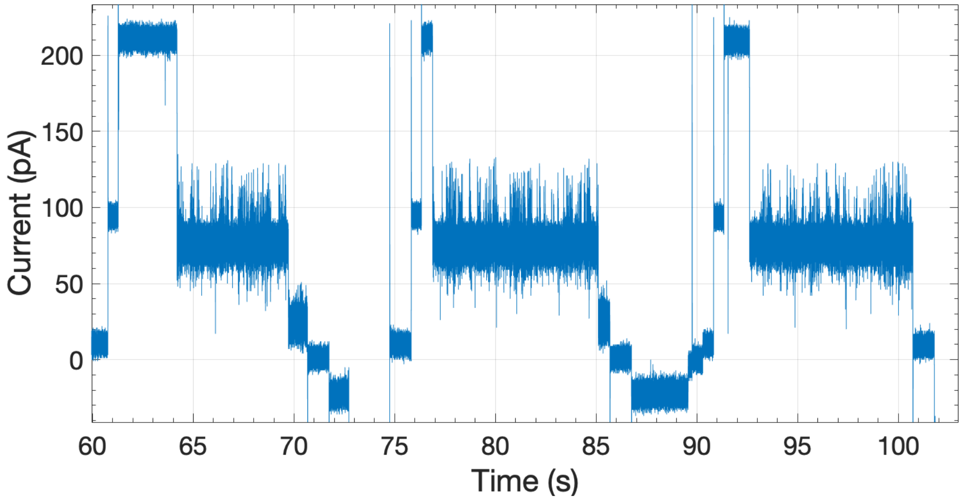


NTER homopolymer E (Glutamate)

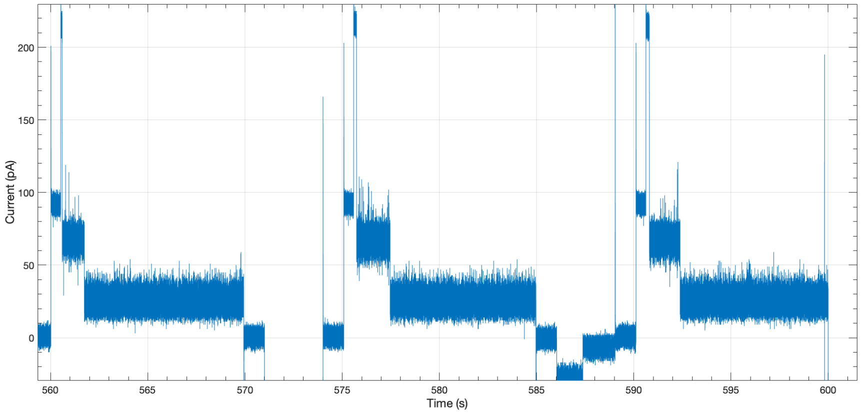


NTER homopolymer Q (Glutamine)

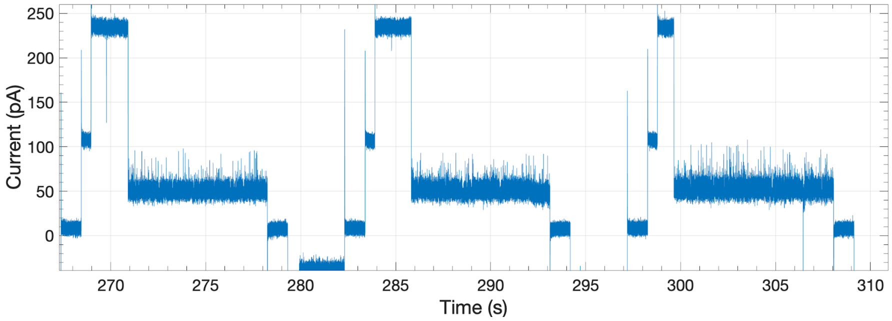

**Supplementary Figure 3 (cont.)**

NTER homopolymer G (Glycine)

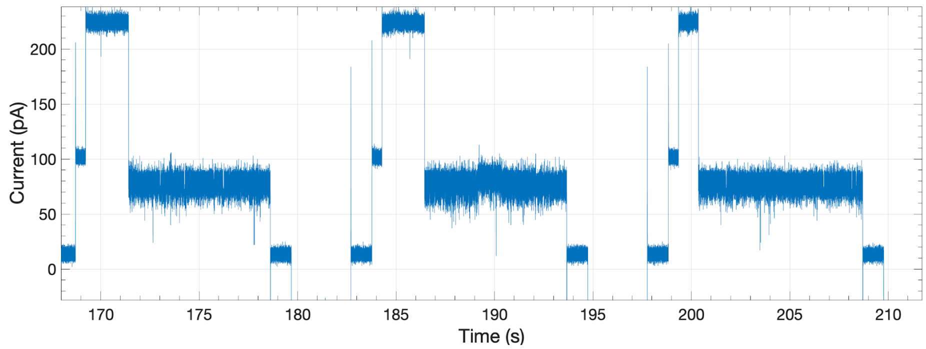


NTER homopolymer H (Histidine)

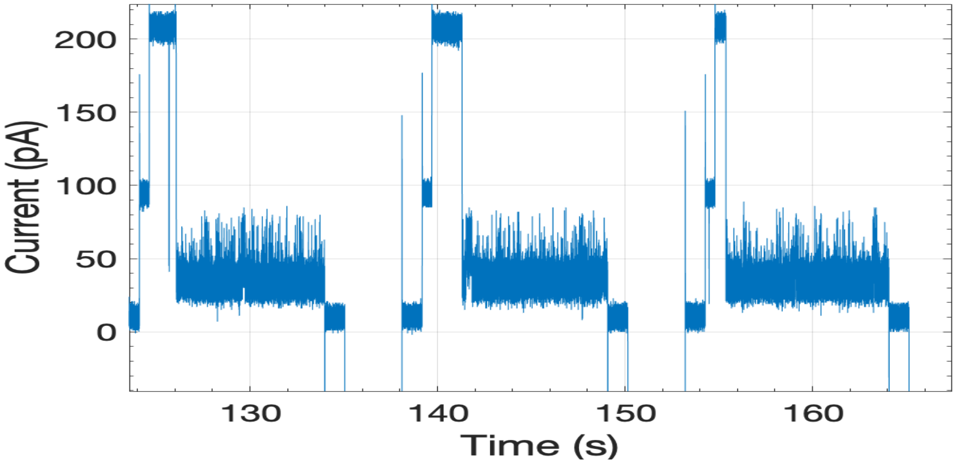


NTER homopolymer M (Methionine)

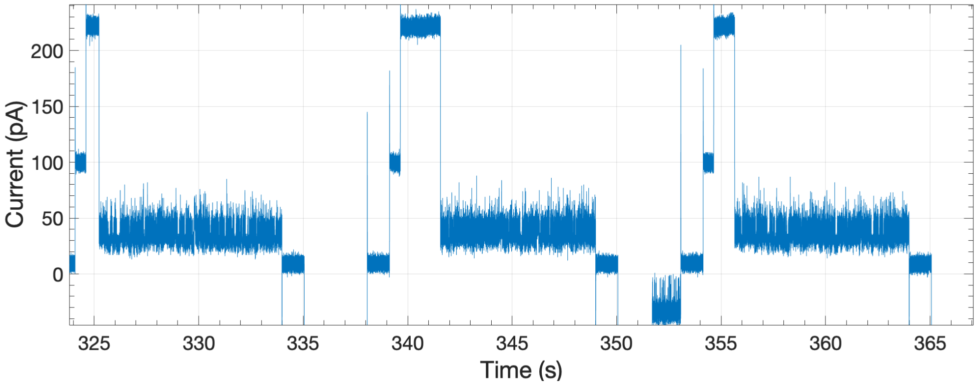

**Supplementary Figure 3  (cont.)**

NTER homopolymer P (Proline)

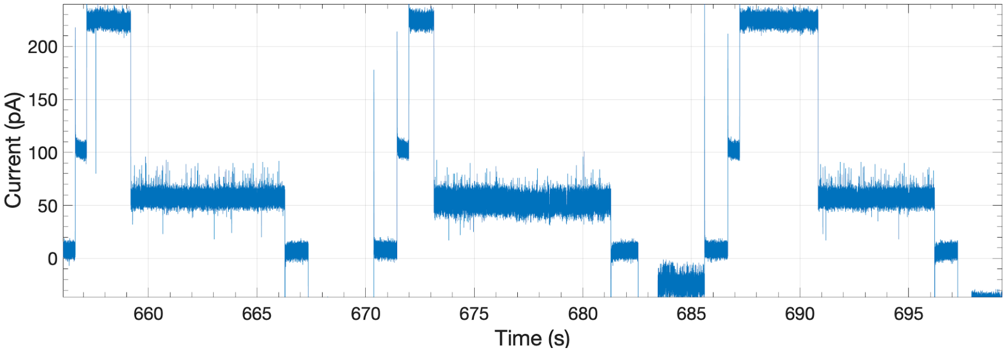


NTER homopolymer S (Serine)

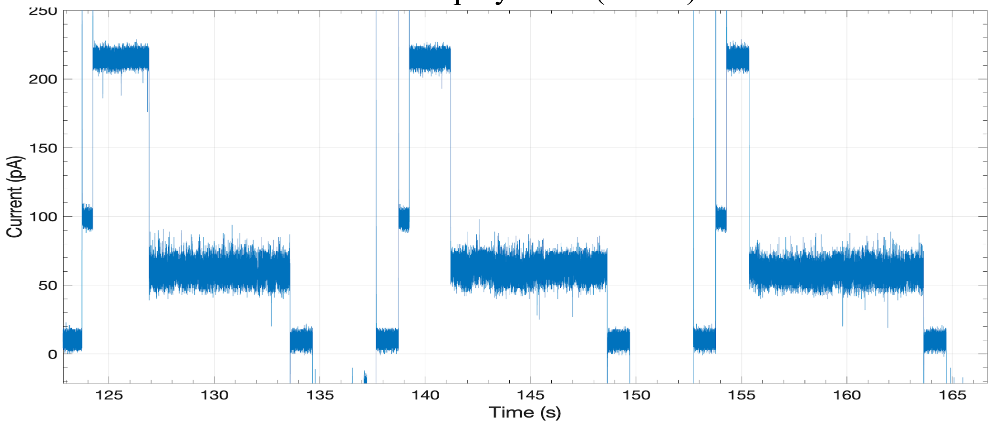


NTER homopolymer T (Threonine)

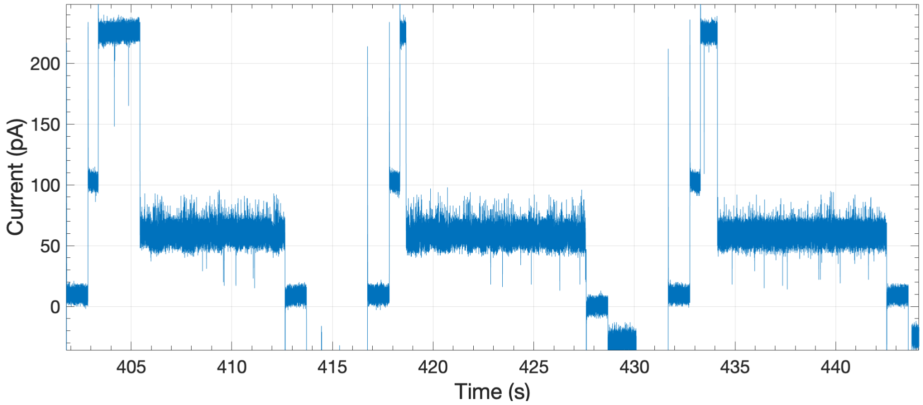

**Supplementary Figure 3 (cont.)**

NTER PKA motif

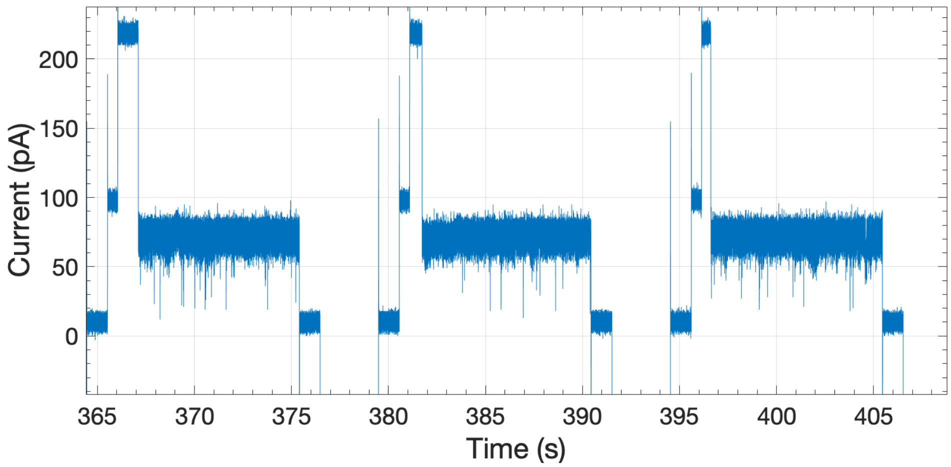


NTER PKA phosphomimetic motif

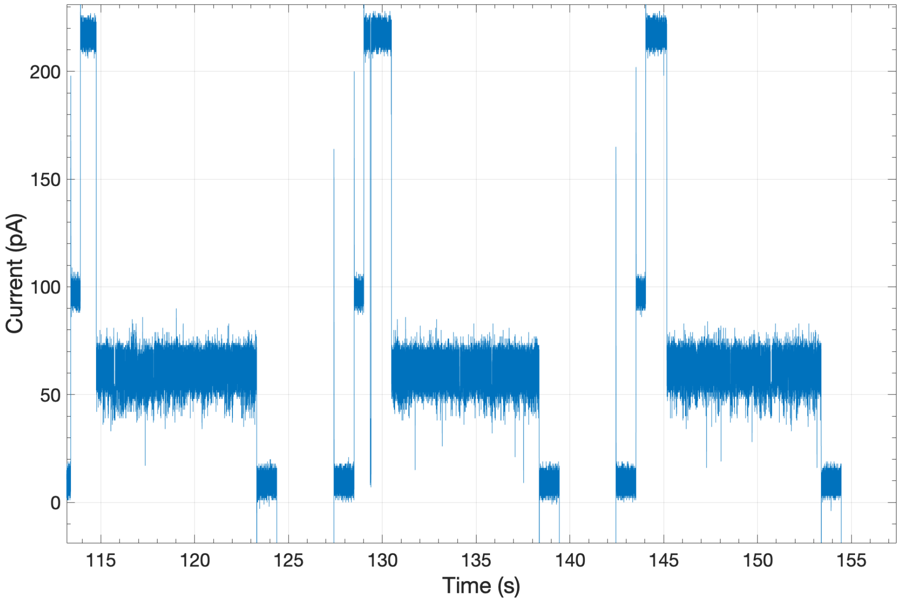

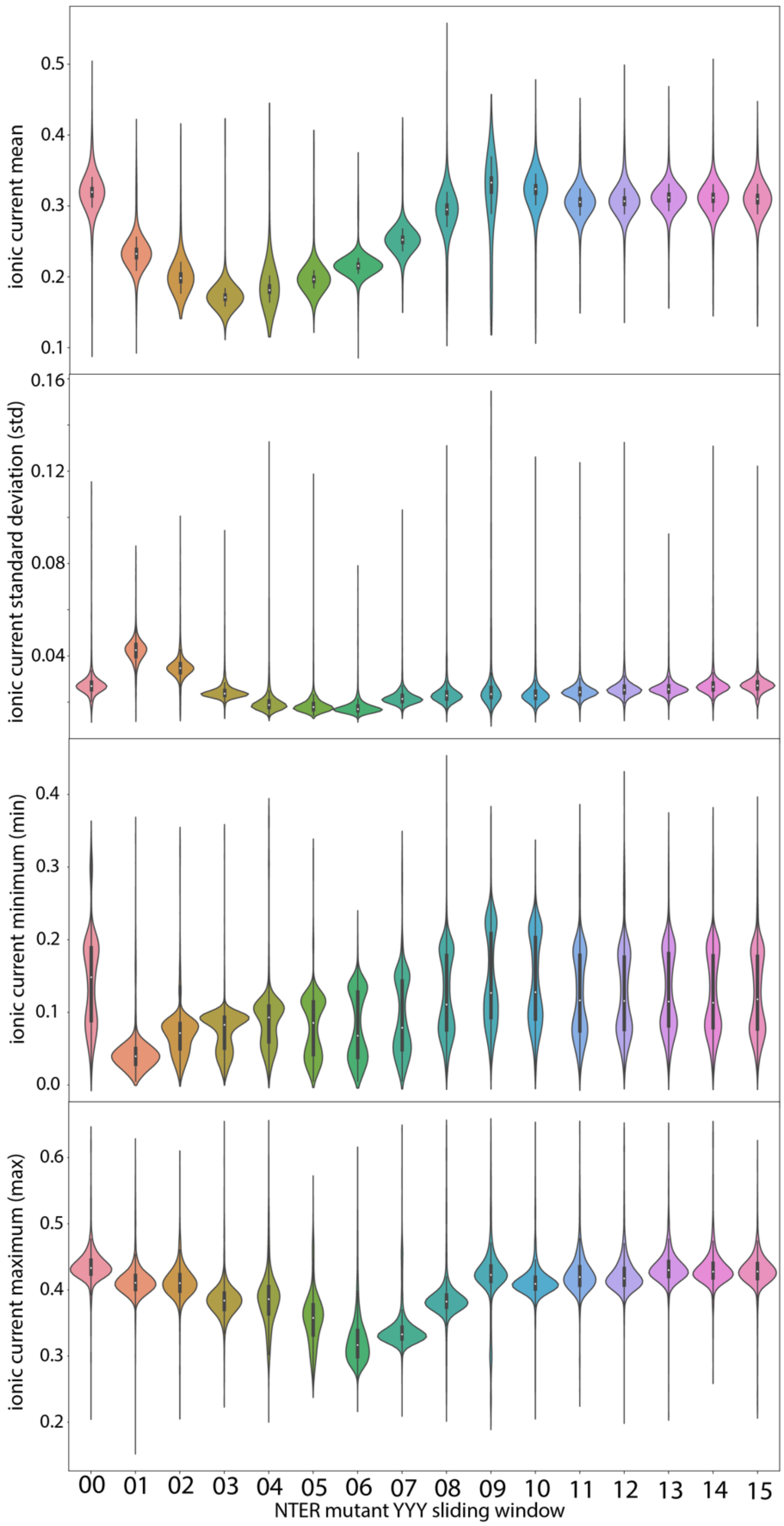


**Supplementary Figure 4** Violin plots showing the ionic current level signal characteristics (mean, std, min, and max. All normalized to the open pore level) of the nanopore capture state for NTERs 00-15. Each NTER distribution is composed of >1000 single-molecule measurements.

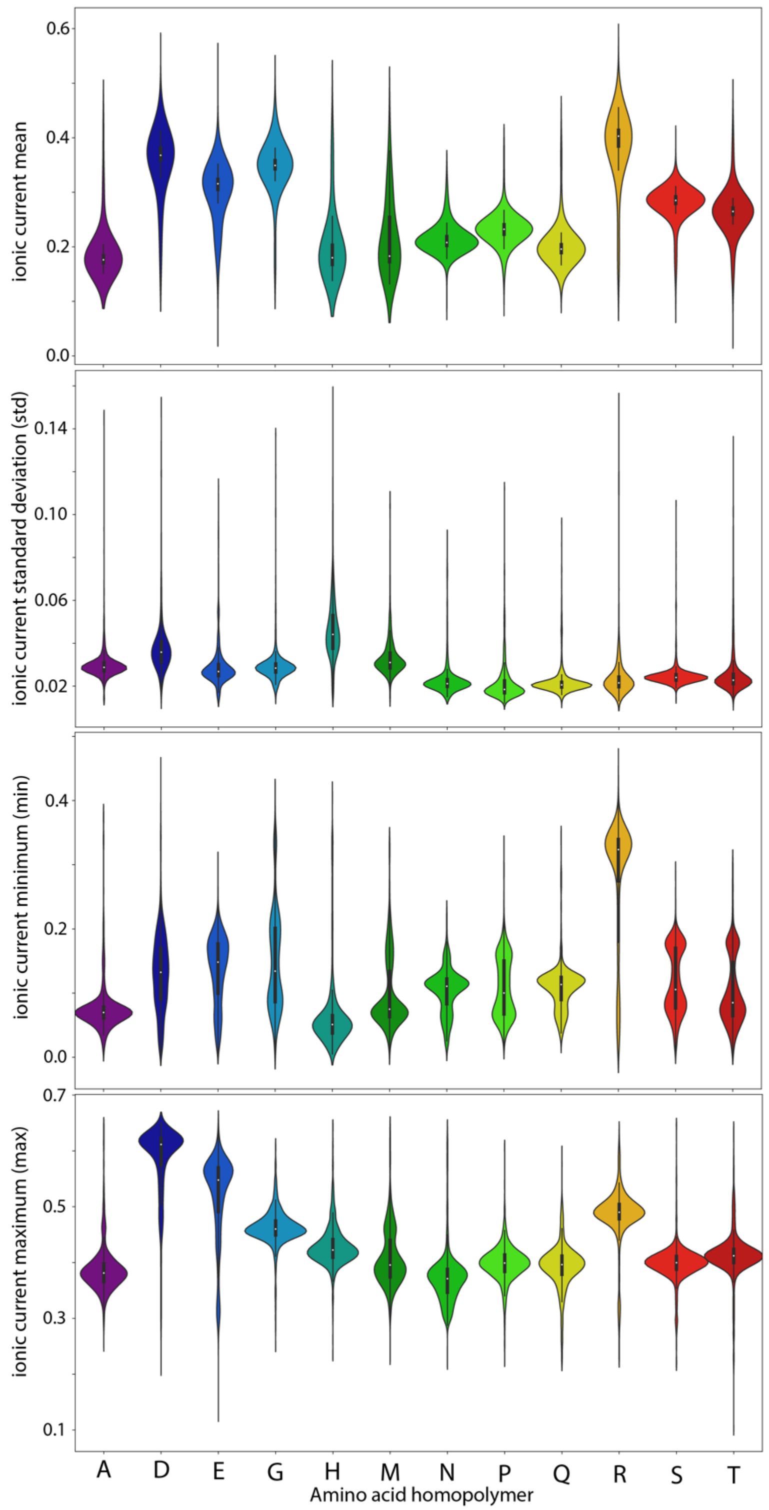


**Supplementary Figure 5** Violin plots showing the ionic current level signal characteristics (mean, std, min, and max. All normalized to the open pore level) of the nanopore capture state for the amino acid homopolymer mutants. Each NTER distribution is composed of >1000 single-molecule measurements.
